# Exosome-LncPICALM-AU1 regulates endothelial–mesenchymal transition in hepatopulmonary syndrome

**DOI:** 10.1101/2020.10.06.327874

**Authors:** Congwen Yang, Yihui Yang, Yang Chen, Jian Huang, Yujie Li, Hongyu Zhi, Xi Tang, Xiaobo Wang, Karine Belguise, Zhengyuan Xia, Jiaoling Ning, Jianteng Gu, Bin Yi, Kaizhi Lu

## Abstract

As important mediators of intercellular communication, exosome have can modulate various cellular functions by transferring a variety of intracellular components to target cells. However, little is known about the role of exosome-mediated communication between distant organs. Hepatopulmonary syndrome (HPS) is a severe lung injury caused by chronic liver disease. A new long noncoding RNA (lncRNA) PICALM-AU1 was found and upregulated in the liver of HPS. It was located in the cholangiocytes of liver and then, secreted as exosome into the serum. PICALM-AU1 carrying serum exosomes induced endothelial-mesenchymal transition (EndMT) of PMVECs and promoted lung injury in vivo and in vitro. Furthermore, overexpression of PICALM-AU1 significantly suppressed miR144-3p and subsequently induced ZEB1 expression. Taken together, our findings identified cholangiocyte-derived exosomal lncRNA PICALM-AU1 plays a critical role in the EndMT of HPS lung. And PICALM-AU1 represents a noninvasive biomarker and potential therapeutic target for HPS.

## Introduction

Hepatopulmonary syndrome (HPS), characterized by hypoxemia and intrapulmonary shunting, occurs in 5–32% of patients with liver disease [1]. HPS significantly increases mortality and worsens functional status and quality of life in patients with cirrhosis [2]. Despite a growing knowledge of the mechanisms involved in the development of HPS, its pathogenesis has not been fully elucidated [3-6]. The core pathogenic feature of HPS includes microvascular changes in pulmonary circulation. Intrapulmonary vascular dilation significantly reduces the efficiency of gas exchange in early-stage HPS lesions. The late stage of HPS is characterized by increased angiogenesis of microvessels that leads to severe hypoxemia and dyspnea [7]. This is a result of the molecules secreted from the damaged liver.

Pulmonary angiogenesis plays a vital role in the development of HPS [8]. Soluble molecules synthesized in the pathological liver, such as the vascular endothelial growth factor, bone morphogenic proteins 2 and 9, placental growth factor, and cyclooxygenase-2, can be transported to the lung via blood, thereby promoting pulmonary microvascularization and aggravating respiratory distress in individuals with HPS [9-12]. Endothelial–mesenchymal transition (EndMT) is characterized by the loss of endothelial cell features and acquisition of specific mesenchymal cell markers that are key in regulating endothelial function and development and structural remodeling of myocardium, blood vessels, and valves [13]. During endothelial dysfunction, EndMT induces vascular remodeling. Numerous studies have implicated EndMT in vascular diseases, including cerebral cavernous malformations, pulmonary hypertension, vascular graft remodeling, and atherosclerosis [14-17]. A lot of work has been done in the early stage, and the intervention of HPS has some effect, but not ideal. Although a recent study has implicated the involvement of exosomes [18], the mechanism(s) underlying the regulation of HPS pathology by EndMT remains to be fully understood.

Researchers have neglected the role of exosomes in the biology of HPS. Exosomes are small extracellular membrane-enclosed vesicles formed by the inward budding of the endosomal membrane and released extracellularly via fusion with the plasma membrane. Exosomal cargos, including noncoding RNAs, proteins and lipids, are implicated in various liver diseases [19, 20]. Exosome function can be divided into two categories. First is for intercellular communication, or short distance communication: exosomes accumulating in the ischemic myocardium are rapidly taken up by infiltrating monocytes to regulate local inflammatory responses [21]. Cancer-derived exosomal miR25-3p promotes the formation of a pre-metastatic niche [22]. Cholangiocyte-derived exosomal lncRNA H19 promotes the activation of hepatic stellate cells and cholestatic liver fibrosis [23, 24]. The second category of exosomes involve cross-organ communication, or long-distance communication. We have previously shown that hepatocyte-derived exosomal miR194 promotes the angiogenesis of pulmonary microvascular endothelial cells (PMVECs) in pulmonary HPS [25]. Exosomes secreted into the serum is transported to the lungs via long-distance transportation, thereby promoting the formation of pulmonary microvessels and aggravation of symptoms associated with HPS. Long noncoding RNAs (lncRNAs) in exosomes also play an important regulatory role for physiological functions and pathological progression, especially in HPS. Thus, we wanted to determine whether exosomes secreted from the liver contain crucial lncRNAs for the regulation of HPS using long-distance communication between organs.

In this study, we have identified a novel lncRNA (MRAK138283, named PICALM-AU1) using microarrays during the screening of the lungs of HPS rats. We have demonstrated that PICALM-AU1 was overexpressed in cholangiocyte-derived exosomes in the liver of HPS rats. Moreover, PICALM-AU1 levels in serum exosomes positively correlated with the severity of lung injury in the rat model of HPS and specimens from patients with HPS. Notably, cholangiocyte-derived exosomal PICALM-AU1 promoted EndMT in PMVECs in the HPS model. Thus, exosome-derived PICALM-AU1 in the liver of HPS rats regulate lung injury via long-distance communication between distant organs. Finally, PICALM-AU1 is a promising candidate for use a non-invasive diagnostic biomarker and therapeutic target for HPS.

## Materials and methods

### HPS patient specimens

All human specimen-related experiments in this study were approved by *Clinical Trials https://clinicaltrials.gov/* (NO. NCT03435406). Patients were diagnosed with HPS based on three parameters: (1) presence of cirrhosis, (2) positive contrast-enhanced echocardiography, and (3) an alveolar-arterial oxygen gradient (P(A-a) O_2_) ≥15 mmHg (or ≥20 mmHg in patients >64 years). Intrapulmonary vascular dilations were assessed using contrast-enhanced echocardiography. Agitated saline causes the formation of >10 μm microbubbles that usually do not pass through the pulmonary capillary bed. Appearance of microbubbles, after injecting into the peripheral vein, first in the right heart, within 3–6 heart actions in the left heart demonstrates abnormal vasodilation of the intrapulmonary capillary bed. Early (<3 heart beats) appearance of microbubbles in the left heart was considered as intracardiac shunting. These patients were excluded from this study since the presence or absence of intrapulmonary shunting could not be judged using contrast-enhanced echocardiography.

### Animal model and treatments

#### Rat model for common bile duct ligation (CBDL)

Rat model of CBDL is a typical model of HPS that was generated using a well-established methodology [26, 27]. All animal experiments were approved by the Animal Care Committee of Third Military Medical University, Chongqing, China (NO. AMUWEC2020457). Male Sprague-Dawley rats (200–220 g, 30 rats/group) were anesthetized using chloral hydrate (Sigma-Aldrich, USA). The control (sham) rats were subjected to isolation of the common bile duct without ligation. The lungs of the animals were dissected and analyzed 1, 3, and 5 wk after surgery. Blood samples were aseptically drawn from the abdominal aorta during laparotomy. A 0.2 ml sample of arterial blood was collected in a heparinized gas capillary tube to measure arterial gas levels. Serum was separated from the blood samples (centrifugation at 2,000×g, 4°C) and used to separate exosomes.

#### Exosome treatment

To analyze the function of HPS-exosomes in the rat lung, rats were randomly divided into four groups (ss-Exo, sham-serum exosome; Hs-Exo, HPS serum exosome; ct-Exo, MIBECs-derived Exo; PO-Exo, PICALM-AU1 OE MIBECs-derived exosome). Exosomes isolated from sham and HPS rat sera and wildtype and PICALM-AU1-overexpressing mouse intrahepatic biliary epithelial cells (MIBECs). We injected rats with 100 μg total protein/100 μL three times and once every other day via the caudal vein.

#### Virus treatment

To analyze the function of PICALM-AU1 in the rat lung, we constructed lentiviral particles containing the sequence for PICALM-AU1 overexpression (OE) and knockdown (KD). The LV-NC, LV-PICALM-OE, and LV-PICALM-KD viruses were injected via the caudal vein (each with 100 µL of 2×10^10^ Tranduction Unites (TU)/ml). After two weeks, the rats were subjected to CBDL. To investigate the function of exosomal PICALM-AU1 in the rat lung, MIBECs were treated with LV-NC and LV-PICALM-OE (each with 10 µL of 1×10^9^ TU/mL). After 72 h, we measured PICALM-AU1 expression followed by isolation of exosomes to infect the rats. To analyze the function of PICALM-AU1 in PMVECs, PMVECs were treated with LV-NC, LV-PICALM-OE, and LV-PICALM-KD (each with 10 µL of 1×10^9^ TU/mL). After 72 h, gene expression, including protein levels, were measured.

### Isolation and characterization of exosomes

Medium culturing human patient serum, rat serum, and MIBECs were collected by centrifugation at 2,000×g for 15 min followed by 16,000×g for 20 min at 4°C. Supernatants were collected and ultracentrifuged at 110,000×g for 70 min. Subsequently, pellets were resuspended in sterile phosphate-buffered saline and purified by centrifugation at 110,000×g for 1 h. The exosomes were resuspended in phosphate-buffered saline, filtered through 0.22 μm filters (Millipore, USA), and stored at -80°C for further analysis.

Transmission electron microscopy (Hitachi HT7700, Japan) was used to characterize the morphology of isolated exosomes. qNano (Izon Science, New Zealand) was used for the size distribution of the isolated exosomes following the manufacturer’s instructions. We used western blotting with the anti-CD63 and anti-CD86 antibodies to analyze the protein markers of exosomes.

### Microarray analysis

Total RNA from the liver of rats in the CBDL operation and sham groups were extracted and reverse transcribed. Double-stranded cDNA was labeled using the Quick Amp Labeling Kit (Agilent Technologies Inc, USA) and hybridized to the Array star Rat 8×60K lncRNA Array, version 2.0. Following the washing steps, the arrays were scanned using the Agilent Scanner G2505B and array images were analyzed using the Agilent Feature Extraction software, version 10.7.3.1. Quantile normalization and subsequent data processing were performed using the GeneSpring GX software, version 11.5.1 (Agilent Technologies Inc, USA). Volcano plot filtering was used to identify significantly different lncRNAs, and the threshold to screen upregulated or downregulated lncRNAs was set at a fold change of >±1.5 and P<0.05.

### Tissue harvest and histology

Liver and lung samples were fixed in 4% phosphate-buffered formaldehyde solution (Klinipath, Belgium), dehydrated, embedded in paraffin, and subjected to hematoxylin and eosin (H.E.) staining, Masson staining, immunohistochemistry, immunofluorescence, and fluorescence *in situ* hybridization (FISH). Table S2 lists all the antibodies used in this study.

#### H.*E. staining*

Rat lung tissues were subjected to H.E. staining as described previously [26].

#### Masson staining

The degree of liver fibrosis was scored using Masson-stained liver sections (5 μm thickness), the METAVIR scoring system (4), and quantitatively analyzed using the Cell^D software (Olympus Imaging Solutions, Germany). Data have been expressed as the mean fibrotic area/field (% ± SE) positively stained using Sirius Red. The final score was represented as the mean of the scores determined by two independent researchers who were blinded to the study samples.

#### Immunohistochemistry

Immunohistochemical staining on lung tissue allowed to quantify protein expression levels. Specific anti-VWF, anti-VE-cadherin and anti-Vimentin were used. Slices that underwent immunostaining with omission of primary antibodies or with IgG were used as negative controls. Paraffin-embedded lung sections (5 μm thickness) were deparaffinized, rehydrated by serial immersion in ethanol, and pretreated with citrate buffer. Non-specific binding sites were blocked via incubation in 3% H_2_O_2_ (Merck, Germany) and BSA respectively. Epitope detection was performed using the ultraView Universal DAB Detection Kit (Dako, Denmark). Counterstaining was performed with hematoxylin.

The vascular density of specimens stained for VWF was measured semi-quantitatively using Cell Software (Olympus, Japan). Results are expressed as mean positively stained area (% ± SE) per field. The number of macrophages per high power field (objective 40×) was counted in 15 randomly selected fields for each mouse, and the mean value of the cell counts in these fields was calculated (mean number of macrophages per field ± SE). All final histological scores are represented as the mean of the scores determined by two independent researchers, who were blinded to the study samples.

#### Immunofluorescence

For immunofluorescent double staining, paraffin-embedded lung sections (5 μm thickness) or cell slides were deparaffinized, rehydrated by serial immersion in ethanol and pretreated with EDTA, followed by incubation in 50 mM NH_4_Cl, 0.1% Triton X-100 and 1% BSA. Anti-VE-cadherin, anti-Vimentin, anti-ZEB1 and anti-ZO1 were used as primary antibodies. Slices that underwent immunostaining with omission of primary antibodies or with IgG were used as negative controls. The binding sites of the primary antibodies were revealed with Alexa Fluor-594 goat anti-rabbit and Alexa Fluor-488 goat anti-mouse secondary antibodies (Invitrogen, USA). Nuclei were stained with 4’, 6-diamidino-2-phenylindole (DAPI) (Life Technologies, USA). Samples were visualized with a fluorescence microscope (Olympus, Japan).

#### FISH (Fluorescence in situ hybridization) combined with fluorescent IHC staining

FISH targeting PICALM-AU1 in rat liver tissue sections was performed using a commercially available RNA scope Multiplex Fluorescent Reagent Kit v2 (Advanced Cell Diagnostics, USA) by following the manufacturer’s instruction. Fluorescent IHC staining target PICALM-AU1 was performed after FISH staining as described in the above section (Histopathology, Masson’s Trichrome staining, and immunohistochemistry). Zeiss LSM 700 confocal laser scanning microscopy were used to visualize FISH results (Carl Zeiss, Germany).

### cDNA synthesis and quantitative polymerase chain reaction (qPCR)

LncRNA, miRNA, and mRNA expression was analyzed using total RNA from tissues and cells and the Applied Biosystems 7000 sequence detection system (Applied Biosystems, UK), SYBR Green, and comparative CT method. Values were reported relative to the endogenous control glyceraldehyde-3-phosphate dehydrogenase. All amplification reactions were performed three times independently. Supplementary Table S1 lists the primer sequences used.

### Western blotting

Protein levels were determined using western blotting of rat lung and PMVEC samples as previously described [28] using specific antibodies (Table S2). Blots were visualized using ECL reagents (DAKO, Denmark), and digital images were acquired using the luminescent image analyzer LAS-4000 (General Electric, UK). β-actin was used for the normalization of quantitative densitometry values.

### Cell culture and in vitro experiments

Rat PMVECs and mouse MIBECs were purchased from American Type Culture Collection (ATCC Cell Biology Collection, USA). Cells were maintained at 37°C in RPMI medium (Gibco, USA) supplemented with 10% fetal bovine serum (Invitrogen, USA). For cell transfection experiments, cells were seeded at 60–70% confluency. Vectors were mixed with Lipofectamine 3000 (Promega, USA), diluted in EGM2, and treated for 24 h as previously described [28]. After 24 h, cells were incubated with miR144-3p mimics/inhibitor or sham/HPS exosomes. Subsequently, luciferase activity was measured using the Dual-Luciferase Reporter Assay System (Promega, USA) and GloMax-Multi Detection System Photometer (Promega, USA).

### Statistical analysis

Results were obtained from at least three independent experiments and expressed as mean±standard deviation. Data were analyzed using two-tailed Student’s *t*-test, one-way analysis of variance with Tukey’s post-hoc test or linear regression using GraphPad Prism software version 8.0 (GraphPad Software Inc., USA). *P*≤ 0.05 was considered statistically significant.

## Results

### lncRNA PICALM-AU1 is overexpressed in the HPS liver

We constructed the rat model of HPS using CBDL to identify a key lncRNA involved in the regulation of the progression of HPS (Fig. 1A). The CBDL rat had cirrhosis, low efficiency of pulmonary gas exchange, and excessive angiogenesis of pulmonary microvessels (Fig. S1A–F). We then performed RNA sequencing to compare the RNA levels in the livers of CBDL and sham rats. After filtering data for long noncoding RNA annotation and expression levels, we identified 88 and 10 lncRNAs that were upregulated and downregulated after CBDL, respectively. We chose the top 4 lncRNAs as primary candidates (Fig. 1B, C). qPCR analysis showed that MRAK138283 was upregulated in the early pathological phase in the liver of CBDL rat (Fig. 1E). BC158594 and MRAK079490 were induced during the late pathological phase (Fig. S1G, H). MRAK144056 did not show a specific trend of expression during the pathological progression of HPS (Fig. S1I). We used the following criteria to choose the lncRNA(s) involved in regulating the progression of HPS: (1) expression in the liver, (2) high expression in the early pathological stage of HPS, and (3) pathophysiological role of secretion from the liver into the lung. Thus, we choose to study MRAK138283 owing to its novelty. MRAK138283 (NCBI: LOC102550036, Ensembl: ENSRNOG00000062120) is encoded by the antisense strand of chromosome 1 in the rat and is upstream of the Picalm gene with two exons spanning 368 bp in the coding sequence (Fig. 1D). Accordingly, we named this lncRNA “PICALM-AU1.”

**Fig. 1.**
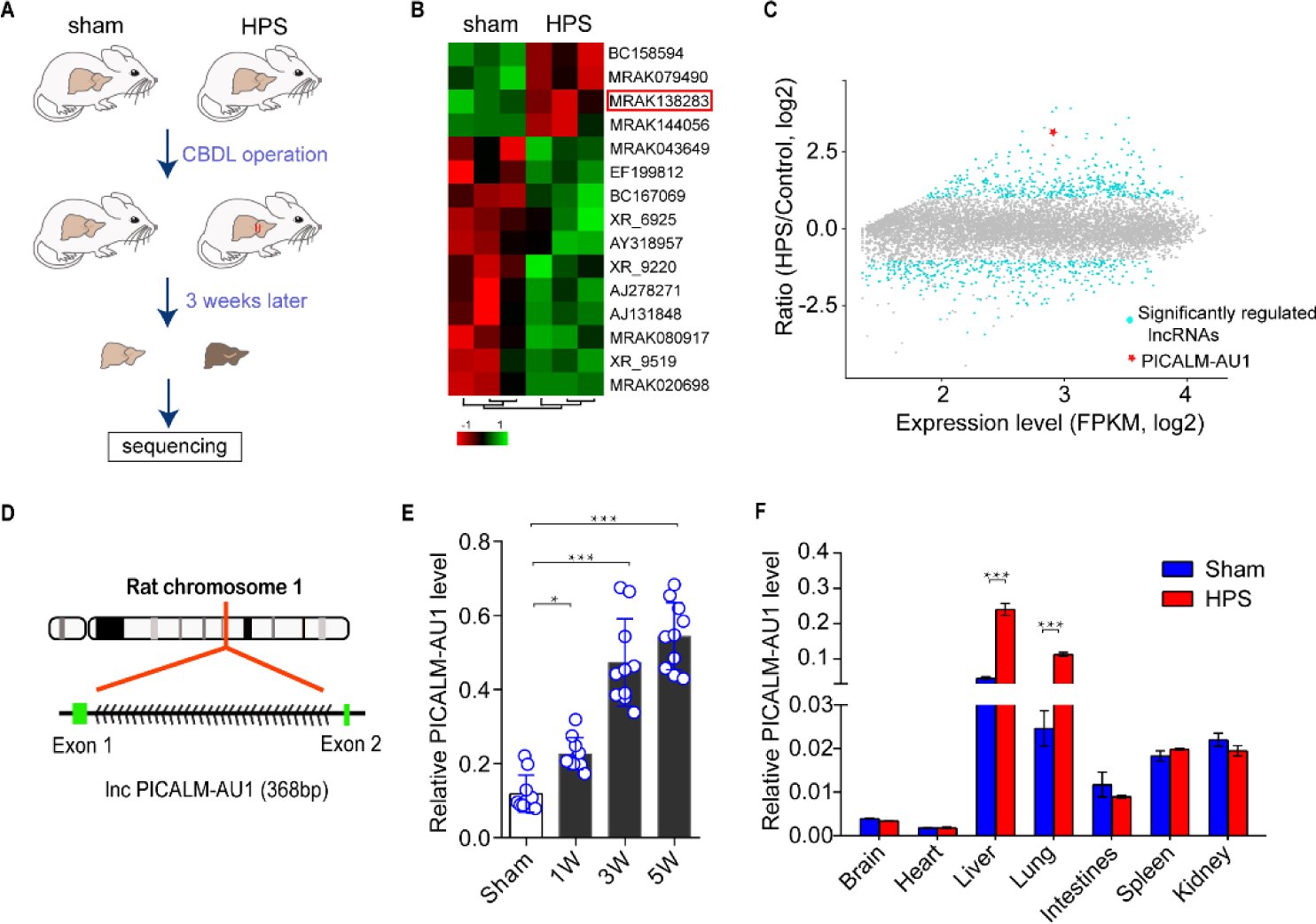
Overexpression of the long noncoding RNA (lncRNA) PICALM-AU1 in the liver of rats with hepatopulmonary syndrome (HPS) A, Strategy involved in generating the rat model of common bile duct ligation (CBDL). B, Analysis of differential gene expression using deep sequencing of the lncRNA array. C, Differentially regulated lncRNAs have been highlighted in light blue (Red, PICALM-AU1). D, The genomic location of PICALM-AU1 in the rat genome. E–F, Quantitative polymerase chain reaction (qPCR) for the expression of PICALM-AU1 across the different stages of HPS in the liver and tissues in the third week of HPS rats. Statistical significance relative to sham group, Student’s *t-*test: *P<0.05, **P<0.01, ***P<0.001, n=10.

Furthermore, we analyzed the expression of PICALM-AU1 in different phases of the liver and normal tissues of sham and CBDL operated rats. PICALM-AU1 was overexpressed in the liver and lung (Fig. 1F) and significantly high in the liver of rats during the first week of HPS (Fig. 1E).

### Overexpression of lncRNA PICALM-AU1 in the liver of HPS rats and its secretion via serum exosomes

To determine the pattern of expression, we first examined PICALM-AU1 levels in the liver of sham and HPS rats using FISH. As expected, PICALM-AU1 was overexpressed in the livers of HPS rats than that in sham rats. The signal from PICALM-AU1 was primarily observed in rat cholangiocytes. Immunofluorescence using CD63, an exosomal surface marker, indicated that CD63 was upregulated in cholangiocytes (Fig. 2A).

**Fig. 2.**
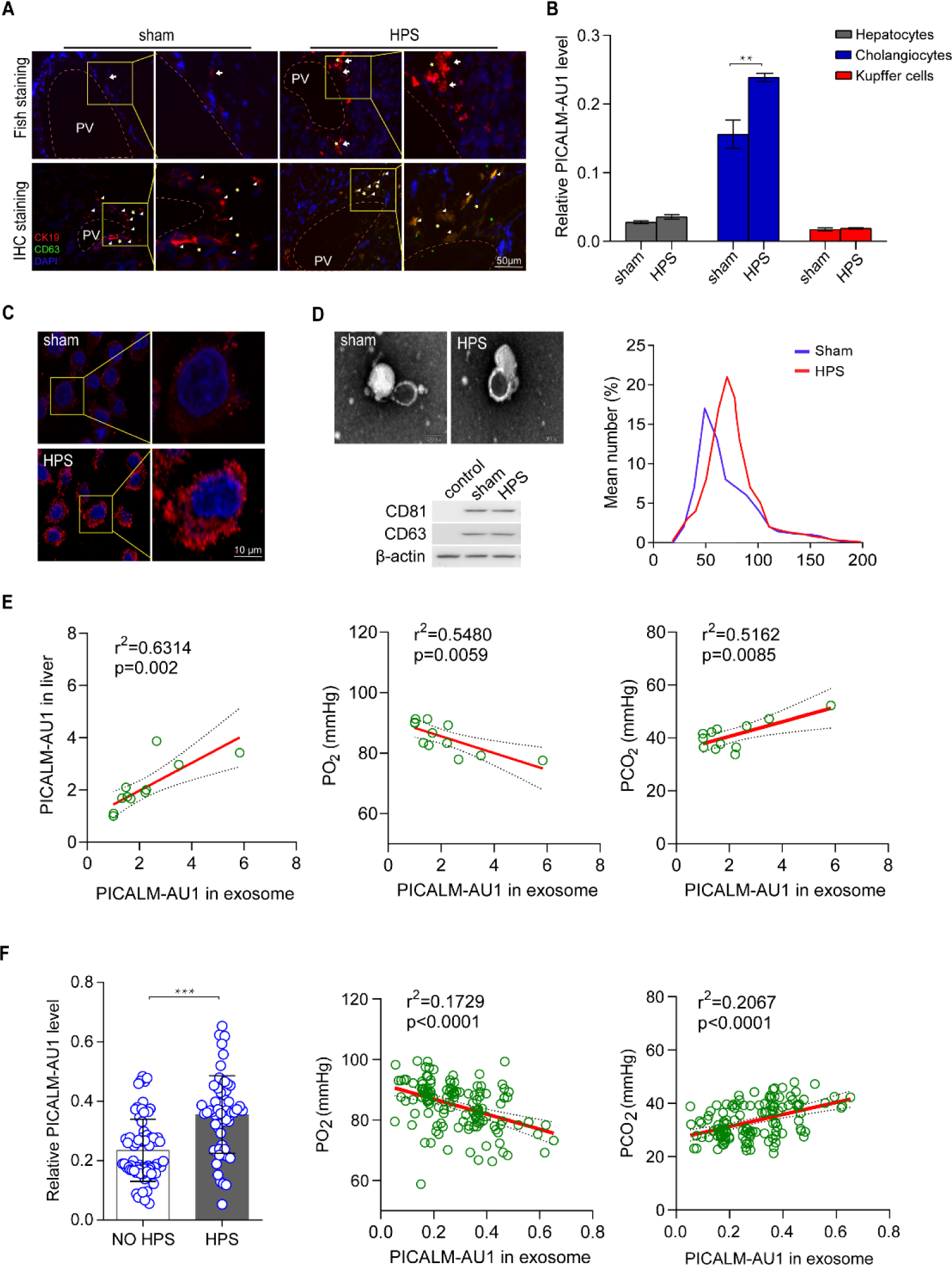
Expression and exosomal secretion of PICALM-AU1 in cholangiocytes. A, Representative images of fluorescence *in situ* hybridization for PICALM-AU1 in the liver (upper panel). Representative images of immunofluorescence for CK19 and CD63 (lower panels). Colocalization of CK19 and CD63 have been indicated using white triangles. Bile duct, yellow *; Portal vein, PV. B, qPCR analysis for the expression of PICALM-AU1 in cholangiocytes, primary hepatocytes, and Kupffer cells. C, FISH for the subcellular localization of PICALM-AU1 in cholangiocytes cell. D, Transmission electron micrographs (upper panel) and western blotting (lower panel) of exosomes isolated from the serum of sham and HPS rats (upper line). The number of exosomes were analyzed (right). E, Correlation between hepatic PICALM-AU1 and serum exosomal PICALM-AU1 (left), partial pressure of oxygen (PO_2_) and exosomal PICALM-AU1 (middle), partial pressure of carbon dioxide (PCO_2_) and exosomal PICALM-AU1 (right). F, Expression of exosomal PICALM-AU1 in 56 and 73 patients with HPS and chronic liver without HPS (left), respectively. We also analyzed the correlation between PO_2_ and exosomal PICALM-AU1 (middle) and PCO_2_ and exosomal PICALM-AU1 (right).

To confirm the overexpression of PICALM-AU1 in the cholangiocytes, we detected the expression of PICALM-AU1 in the three main types of liver cells (cholangiocytes, Kupffer cells, and hepatocytes). As shown in Fig. 2B, the mRNA levels of PICALM-AU1 were high in cholangiocytes of the CBDL rats and were associated with the progression of HPS. PICALM-AU1 primarily localized to the cytoplasm (with slight localization in the nucleus) of cholangiocytes in the livers of HPS rats as compared to its localization in the sham rats (Fig. 2C).

Based on these findings, we speculated that PICALM-AU1 may be secreted from cholangiocytes in exosomes and functions in the lung. We isolated exosomes from sham and HPS rats and measured the expression of PICALM-AU1 (Fig. 2D). Correlation analysis showed that the mRNA levels of hepatic PICALM-AU1 positively correlated with the PICALM-AU1 content in exosomes (r=0.7946; p=0.002) and partial pressure of carbon dioxide (PCO_2_; r=0.7185; p=0.0085). The mRNA levels of hepatic PICALM-AU1 negatively correlated with the partial pressure of oxygen (PO_2_; r=0.7403; p=0.0059, Fig. 2E).

To confirm these data, we selected 56 patients with HPS from 135 patients with chronic cirrhosis (Fig. 2F and Table 1). Patients with HPS had vertical dyspnea, positive type-B ultrasound, higher arterial PCO_2_, and lower arterial PO_2_ than those in patients without HPS. Serum levels of Exo-PICALM-AU1 were higher in patients with HPS as compared to those in patients without HPS. Serum exosomal levels of PICALM-AU1 negatively correlated with PO_2_, but positively correlated with PCO_2_. These results indicate that cholangiocytes are the primary source of serum exosomal PICALM-AU1 and PICALM-AU1 levels were associated with pathological progression of HPS.

### Exo-PICALM-AU1 promotes PMVECs EndMT in the lungs of rats

The cellular mechanism involved in EndMT, involved in various cardiovascular pathologies, includes tissue fibrosis after injury. EndMT in PMVECs plays an important role in regulating angiogenesis and blood vessel remodeling [29]. Immunohistochemistry and western blotting showed a decrease and induction of the expression of VE-cadherin (endothelial biomarker) and vimentin (mesenchymal cell biomarker), respectively, in the lungs of HPS rats (Fig. S2A, B). Correlation analysis showed that VE-cadherin levels negatively correlated with that of vimentin in the lungs of HPS rats (r=0.8259; p<0.0001; Fig. S2A). We also detected the expression of PICALM-AU1 in the lungs with HPS. This indicated the induction in the expression of PICALM-AU1 in the 3 w and 5 w marks in the lungs of HPS rats (Fig. S2C).

To identify whether Exo-PICALM-AU1 stimulated EndMT in PMVECs, we used HPS rat-derived exosomes to treat normal rats (Fig. 3A). Using sham-serum Exo treatment as the control, we observed the decrease in mRNA and protein levels of VE-cadherin in the HPS rat-derived exosome treatment group. These samples exhibited increased mRNA and protein levels of vimentin. This was also observed in lungs treated with exosomes derived from PICALM-AU1-overexpressing MIBECs (Fig. 3B, C). The mRNA levels of PICALM-AU1 also increased dramatically in the lungs treated with HPS exosomes and MIBEC-derived exosomes that overexpressed PICALM-AU1 (Fig. 3B). Correlation analysis showed that PICALM-AU1 levels negatively correlated with that of VE-cadherin, and positively correlated with that of vimentin in the lungs of HPS rats (r_VE-cadherin_=0.9572, p<0.0001; r_Vimentin_=0.9813, p<0.0001; Fig. 3C).

**Fig. 3.**
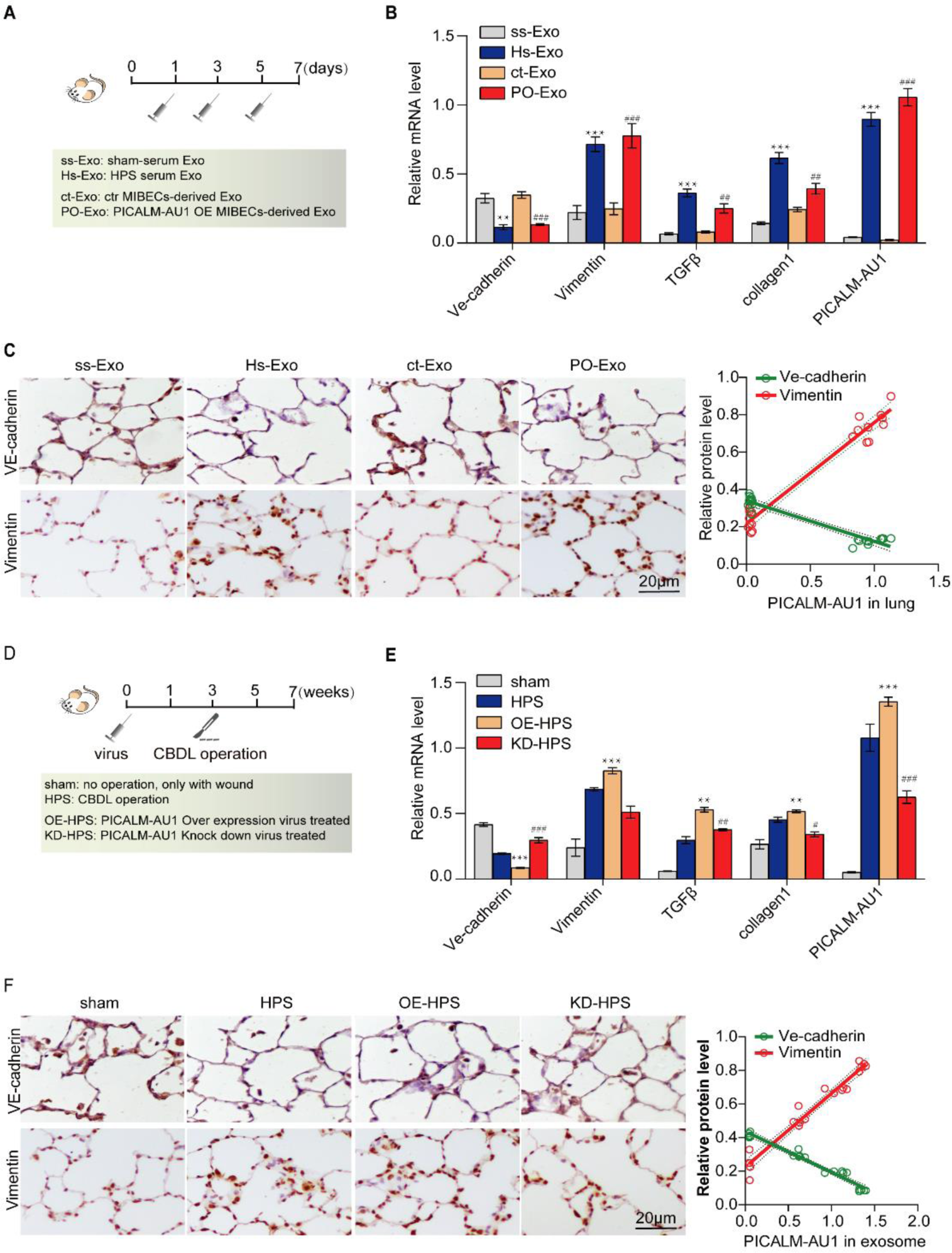
Exo-PICALM-AU1 promoted endothelial–mesenchymal transition (EndMT) in pulmonary microvascular endothelial cells (PMVECs) A, Experimental protocol used for treating rats with exosomes. B, qPCR analysis for relative mRNA levels. Statistical significance relative to ss-Exo treated group, *P<0.05, **P<0.01, ***P<0.001; relative to ct-Exo treated group, #P<0.05, ##P<0.01, ###P<0.001, n=5. C, Immunohistochemistry showing the downregulation of VE-cadherin (endothelial cell marker) and upregulation of vimentin (mesenchymal cell marker) in rats with progressing HPS; (ss-Exo, Serum exosomes from sham rat; HPS-Exo, serum exosomes from 3-wk-old CBDL rats; ct-Exo, exosomes from the culture medium of control mouse intrahepatic biliary epithelial cells [MIBECs]; PO-Exo, exosomes from PICALM-AU1-overexpressing MIBECs). (Left, Immunohistochemistry; Right, Linear regression analysis of VE-cadherin and vimentin). D, Experimental set-up for the overexpression and knockdown of PICALM-AU1 in HPS rats. E, qPCR analysis for relative mRNA levels. Statistical significance relative to the sham group, *P<0.05, **P<0.01, ***P<0.001; relative to the HPS group, #P<0.05, ##P<0.01, ###P<0.001, n=5. Data were compared using two-way analysis of variance. F, Immunohistochemistry for the downregulation of VE-cadherin and overexpression of vimentin during HPS in the lungs of rats. (Left, Immunohistochemistry; Right, Linear regression analysis of VE-cadherin and vimentin).

Next, we wanted to investigate whether EndMT in PMVECs was induced by PICALM-AU1. CBDL rats were transfected with lentiviral particles containing the constructs for PICALM-AU1 overexpression or knockdown (Fig. 3D). PICALM-AU1-overexpressing rats manifested with increased pathological changes, downregulated VE-cadherin, and upregulated vimentin than those in rats of the sham and HPS group. PICALM-AU1 knockdown partially reversed the changes in the pathology of HPS and expression of VE-cadherin and vimentin (Fig. 3E, F). Correlation analysis showed that PICALM-AU1 levels negatively correlated with that of VE-cadherin, and positively correlated with that of vimentin in the lungs of HPS rats (r_VE-cadherin_=0.9816, p<0.0001; r_Vimentin_=0.9793, p<0.0001; Fig. 3F).

We used exosomes from HPS rat sera and PICALM-AU1-overexpressing MIBECs to treat PMVECs and determine the effect of Exo-PICALM-AU1 on EndMT in PMVECs. The exosomal content from the HPS rats and MIBECs stimulated the proliferation and migration of PMVECs (Fig. 4A–C). Depletion of PICALM-AU1 suppressed the proliferation and migration of PMVECs (Fig. 4D–F). Thus, PICALM-AU1 induced EndMT in PMVECs and stimulated the pathological progression of HPS.

**Fig. 4.**
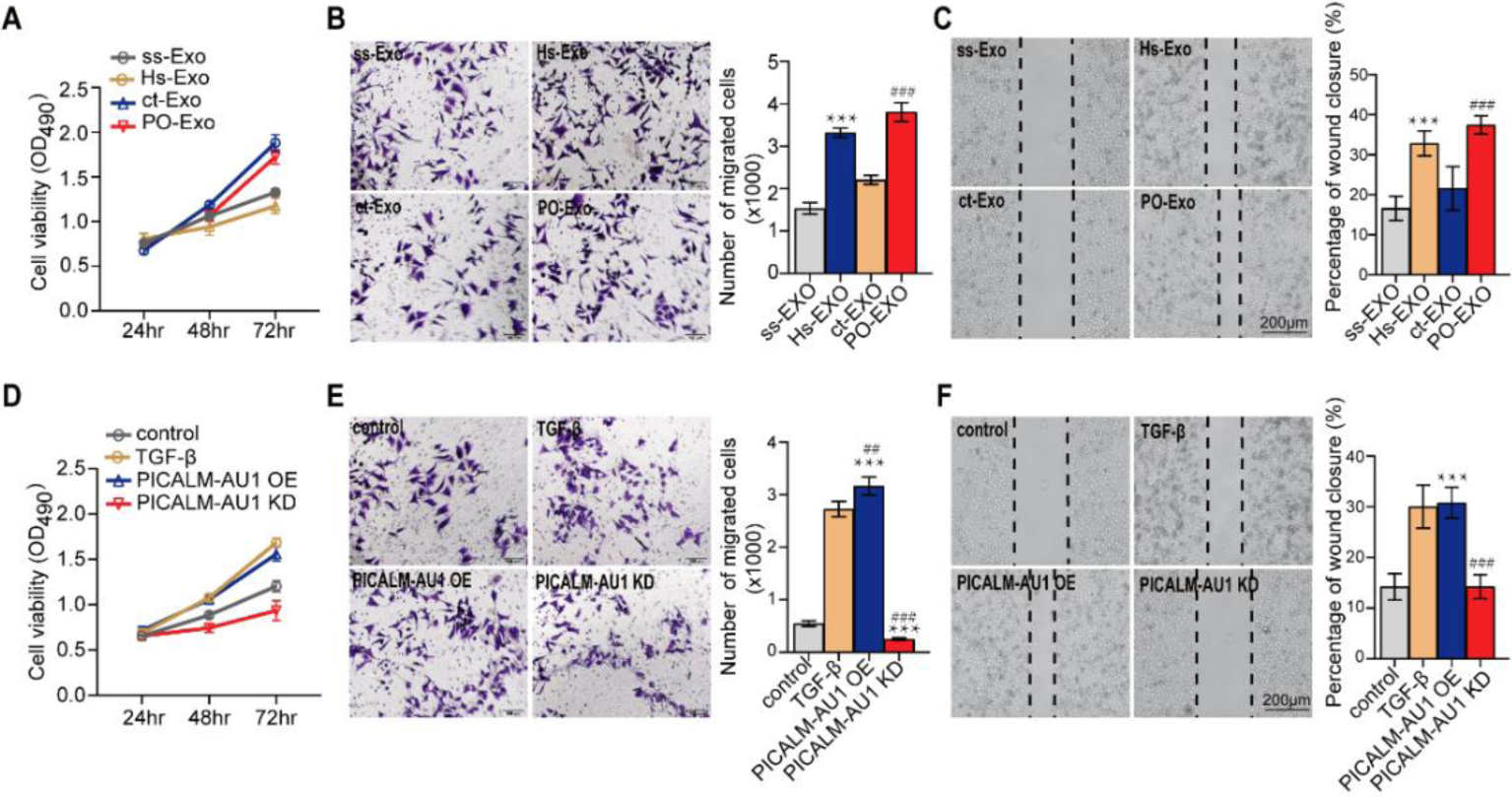
Exo-PICALM-AU1 promoted PMVEC proliferation and migration. A–C, Exosome-treated PMVECs. HPS rat serum-derived exosomes and PICALM-AU1-over expression MIBECs cells derived exosome induced PMVEC proliferation (A) and migration (B, C). Statistical significance relative to the ss-Exo treated group, *P<0.05, **P<0.01, ***P<0.001; relative to the ct-Exo treated group, #P<0.05, ##P<0.01, ###P<0.001, n=5. D–F, Lentivirus-mediated over expression and knockdown of PICALM-AU1 in PMVECs. D, Overexpression of PICALM-AU1 induced PMVEC proliferation (D) and migration (E, F). Depletion of PICALM-AU1 reduced PMVEC proliferation and migration; statistical significance relative to control, *P<0.05, **P<0.01, ***P<0.001; relative to TGF-β, #P<0.05, ##P<0.01, ###P<0.001, n=5. Data were analyzed using two-way analysis of variance.

### miR144-3p is a target of PICALM-AU1

To investigate how PICALM-AU1 regulated EndMT in PMVECs, we analyzed the gene expression network in the lungs of HPS rats using a microarray (data unpublished). microRNA 144-3p was found to be a putative target of PICALM-AU1 (Fig. S3A). PICALM-AU1 contains a binding site for miR144-3p (Fig.5C). We have previously shown that miR144-3p inhibits PMVEC proliferation in the lungs of HPS rats [26], indicating that PICALM-AU1 may regulate EndMT in PMVECs via miR144-3p.

Lungs of HPS rats showed the downregulation of miR144-3p (Fig. S3B). Correlation analysis showed that miR144-3p levels in the lungs of HPS rats negatively correlated with Exo-PICALM-AU1 in the serum exosomes of HPS rats (r=0.9088, p<0.0001; Fig. S4A). miR144-3p levels positively correlated with that of VE-cadherin and negatively correlated with that of vimentin (r_VE-cadherin_=0.9523, p<0.0001; r_Vimentin_=0.9558, p<0.0001; Fig. S4D).

miR144-3p in the rat lung was downregulated by HPS exosome treatment and PICALM-AU1-overexpressing MIBEC-derived exosomes (Fig. S3C). Correlation analysis showed that miR144-3p levels in the HPS rat lung negatively correlated with that of PICALM-AU1 in HPS rats (r=0.9017, p<0.0001; Fig. S4B). miR144-3p levels in the HPS rat lung positively correlated with that of VE-cadherin, and negatively correlated with that of vimentin (r_VE-cadherin_=0.9305, p<0.0001; r_Vimentin_=0.895, p<0.0001; Fig. S4E).

qPCR showed decreased levels of miR144-3p upon the overexpression of PICALM-AU1 than that in the control HPS group. Depletion of PICALM-AU1 partially restored miR144-3p levels (Fig. S3D). Correlation analysis showed that miR144-3p levels in the HPS rat lung negatively correlated with that of PICALM-AU1 (r=0.9658, p<0.0001; Fig. S4C). miR144-3p levels in the HPS rat lung positively correlated with that of VE-cadherin, and negatively correlated with that of vimentin (r_VE-cadherin_=0.9809, p<0.0001; r_Vimentin_=0.9512, p<0.0001; Fig. S4F). Thus, miR144-3p levels in the HPS rat lung negatively correlated with that of PICALM-AU1 and EndMT in PMVECs.

### PICALM-AU1 suppresses the expression of miR144-3p

First, to identify whether miR144-3p can regulate EndMT in PMVECs, we treated PMVECs with miR144-3p mimics and inhibitor. The overexpression of miR144-3p stimulated EndMT by upregulating and downregulating vimentin and VE-cadherin, respectively. Downregulation of miR144-3p inhibited EndMT by suppressing the expression of vimentin (Fig. 5A). Correlation analysis showed that miR144-3p levels in PMVECs positively correlated with that of VE-cadherin, and negatively correlated with that of vimentin (r_VE-cadherin_=0.9525, p<0.0001; r_Vimentin_=0.9305, p<0.0001; Fig. 6B).

**Fig. 5.**
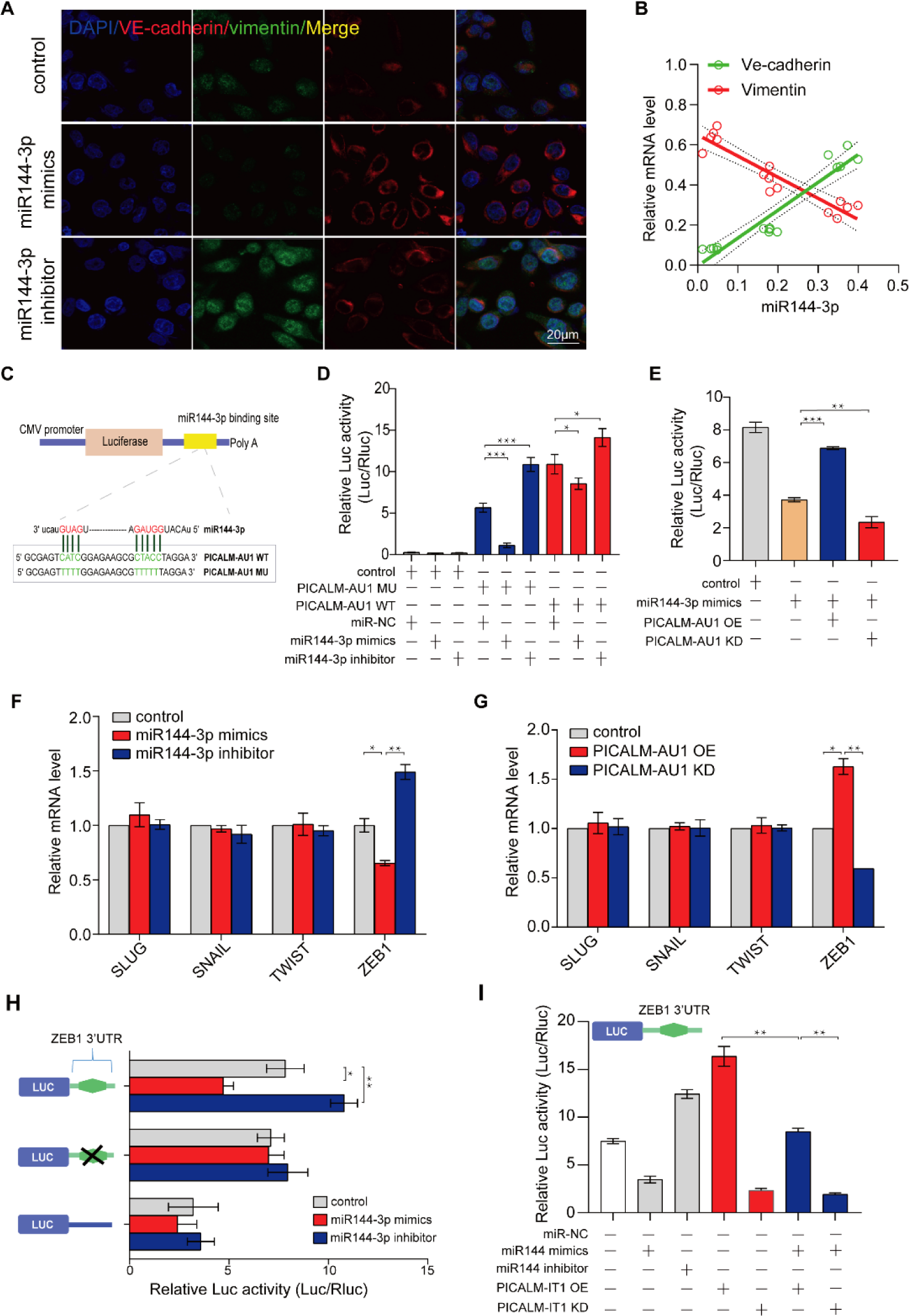
PICALM-AU1 promoted EndMT in PMVECs by inhibiting miR144-3p/Zeb1. A, Immunofluorescence for the expression of VE-cadherin and vimentin in PMVECs treated with miR144-3p mimics and inhibitor. B, Linear regression analysis of miR144-3p levels and VE-cadherin/vimentin levels in PMVECs treated with miR144-3p-specific mimics and inhibitor. C, Schematic for the predicted binding sites of miR144-3p on lncRNA PICALM-AU1. D, Luciferase activity of psiCHECK2-PICALM-AU1 and psiCHECK2-PICALM-AU1mut in PMVECs transfected with the indicated miRNA mimics (n=3). psiCHECK2-miR144-3p (3×) was used as the positive control. Data have represented as the ratio of Renilla luciferase activity to Firefly luciferase activity. E, Luciferase activity of psiCHECK2-PICALM-AU1 in PICALM-AU1-overexpressing or -depleted PMVECs. F, qPCR analysis for the expression of key EndMT-associated transcription factor in PMVECs overexpressing/depleted of miR144-3p. G, qPCR analysis for the expression of key EndMT-associated transcription factor in PMVECs overexpressing/depleted of PICALM-AU1. H, Dual-luciferase assay for the inhibition of ZEB1 by miR144-3p. I, Dual-luciferase assay for the regulated expression of miR144-3p/ZEB1 by PICALM-AU1.

**Fig. 6.**
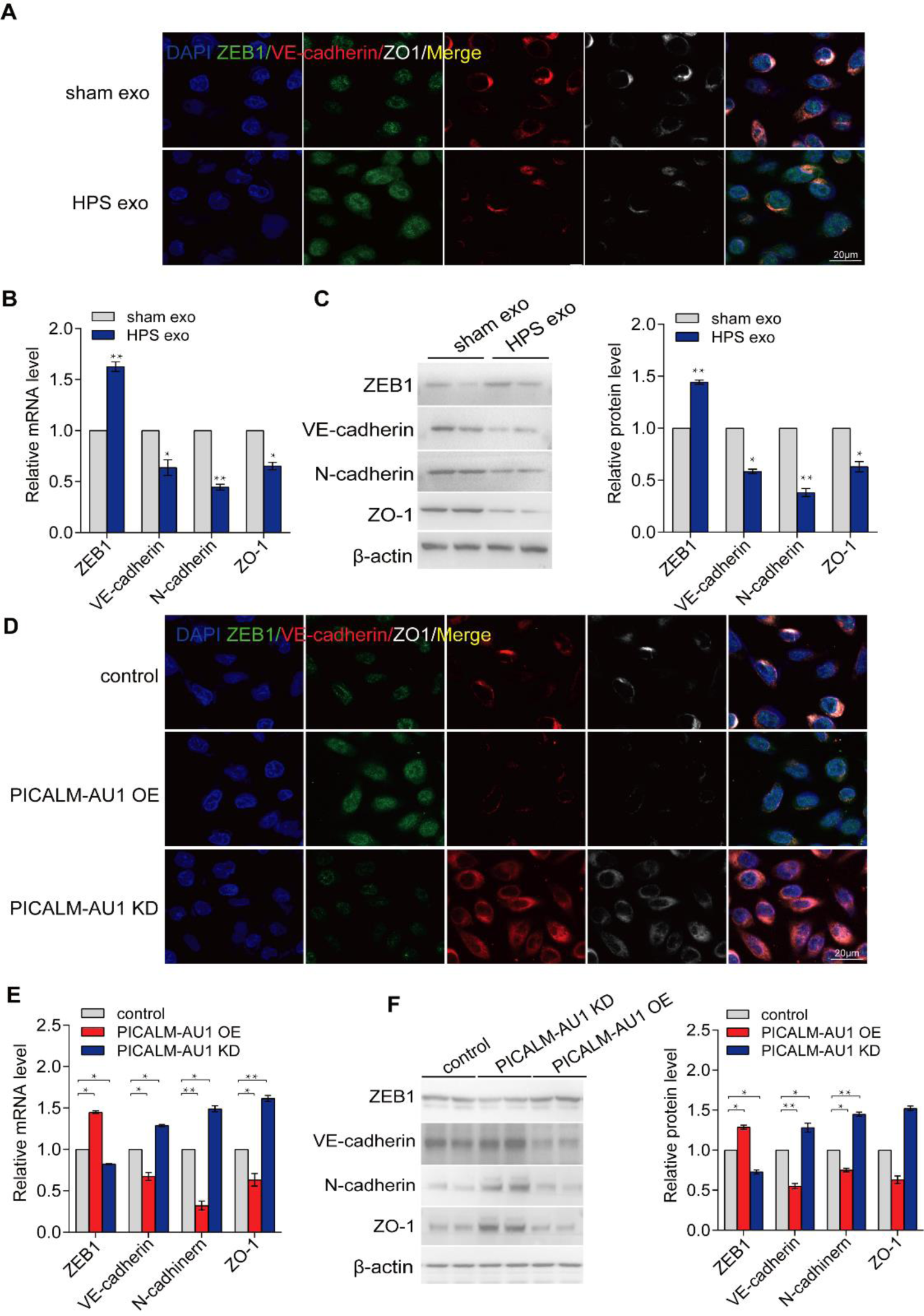
PICALM-AU1 induced EndMT in PMVECs by inhibiting miR144-3p/ZEB1. A–C, Immunofluorescence, qPCR, and western blotting showing the overexpression of ZEB1 and downregulation of VE-cadherin and ZO-1 after treatment with HPS exosomes as compared to those after treatment with sham exosomes. D–F, Immunofluorescence, qPCR, and western blotting showed the overexpression of ZEB1 and downregulation of VE-cadherin and ZO-1 in PICALM-AU1-overexpressing samples. Depletion of PICALM-AU1 reduced ZEB1 expression and induced the expression of VE-cadherin and ZO-1.

We used a luciferase reporter system to analyze the regulatory effect of PICALM-AU1 on miR144-3p (Fig. 5C). The pGL3 vector containing the 3′ untranslated region (UTR) of Tie2 (with miR144-3p binding sites **[26]**) downstream of the LUC gene was transfected into PMVECs. Subsequently, miR144-3p mimics and inhibitor were used to treat PMVECs to upregulate and downregulate miR144-3p, respectively. The nuclear fragment of PICALM-AU WT and PICALM-AU1 MUT were used to overexpress PICALM-AU1. Luc activity was reduced by 20% and enhanced to 150% in PICALM-AU1 MUT-transfected PMVECs containing the miR144-3p mimics and inhibitor, respectively. Transfection of the wildtype PICALM-AU1 nuclear fragment into PMVECs restored the Luc activity in cells with the miR144-3p mimics. And the Luc activity reached the maximum in miR144-3p inhibitor treated PMVECs (Fig. 5D). Thus, PICALM-AU1 negatively regulated the expression of miR144-3p in PMVECs.

To confirm this, we used lentiviral constructs to overexpress and knockdown PICALM-AU1 in PMVECs. PICALM-AU1 overexpression enhanced Luc activity by 1.6-fold in PMVECs. PICALM-AU1 knockdown reduced Luc activity to 25% (Fig. 5E). These results suggested that PICALM-AU1 can regulate EndMT in PMVECs by inhibiting the expression of miR144-3p.

### miR144-3p inhibits EndMT in PMVECs via the ZEB1 transcription factor

To analyze the role of miR144-3p in regulating EndMT in PMVECs, we used miRWalk, miRtarBase, and TargetScanHuman (http://mirwalk.umm.uni-heidelberg.de/, http://mirtarbase.mbc.nctu.edu.tw/php/index.php, and http://www.targetscan.org/vert_71/) to determine the targets of miR144-3p. ZEB1 was a putative target of miR144-3p (Table S3). ZEB1, Zinc Finger E-Box Binding Homeobox 1, acts as a transcriptional repressor for interleukin-2 (IL-2) [30]. It also binds to and suppresses the transcription of E-cadherin to induce epithelial– mesenchymal transition by recruiting SMARCA4/BRG1 [30-32]. Thus, we overexpressed and knocked down miR144-3p using specific mimics and inhibitor, and used qPCR to determine the expression of ZEB1, SNAIL, TWIST, and SLUG. miR144-3p mimics reduced the mRNA levels of ZEB1 in PMVECs, while the miR144-3p inhibitor upregulated ZEB1 (Fig. 5F). Next, we overexpressed or knocked down PICALM-AU1 in PMVECs. Overexpression of PICALM-AU1 upregulated ZEB1 in PMVECs, whereas the depletion of PICALM-AU1 downregulated ZEB1 (Fig. 5G). We then used dual-luciferase assays to analyze miR144-3p-mediated regulation of ZEB1. ZEB1 3′ UTR was transfected into PMVECs and the cells were treated with miR144-3p mimics or inhibitor. miR144-3p bound to the 3′ UTR of ZEB1 and inhibited Luc activity. Mutating the binding site abrogated the potential of miR144-3p to inhibit Luc activity (Fig. 5H). We overexpressed PICALM-AU1 in PMVECs; this partially recovered Luc activity by inhibiting miR144-3p (Fig. 5I). These results showed that miR144-3p may inhibit EndMT in PMVECs via ZEB1, and PICALM-AU1 may stimulate EndMT by inhibiting miR144-3p.

### PICALM-AU1 promotes EndMT in PMVECs via the miR144-3p/ZEB1 axis

To confirm whether PICALM-AU1 regulates EndMT in PMVECs via the miR144-3p/ZEB1 axis, we treated PMVECs with sham and HPS rat serum exosomes. Immunofluorescence, qPCR, and western blotting showed the upregulation of ZEB1 along with downregulation of VE-cadherin, N-cadherin, and ZO-1 after HPS Exo treatment (Fig. 6A–C). Similarly, lentivirus-mediated overexpression of PICALM-AU1 upregulated ZEB1 and downregulated VE-cadherin and ZO-1. Depletion of PICALM-AU1 downregulated ZEB1 and upregulated VE-cadherin and ZO-1 (Fig. 6D–F). Taken together, these findings suggest that PICALM-AU1 secreted in exosomes from cholangiocytes function in the HPS lung in promoting EndMT in PMVECs via the miR144-3p/ZEB1 regulatory axis.

## Discussion

Previous studies have focused on understanding the underlying mechanism of pulmonary microvascular remodeling and concomitant improvement of pathology associated with HPS [33-38]. However, microvascular remodeling induces a limited effect. Thus, we speculated that liver disease precedes the development of HPS. To that extent, liver secretions may provide important information about the pathology of HPS. We have previously studied the regulation of c-kit^+^ cells in the lung by serum SDF1 that is involved in angiogenesis of the pulmonary microvasculature [4]. In this study, we identified a novel lncRNA, PICALM-AU1, that was primarily expressed in cholangiocytes and secreted in exosomes. Cholangiocyte-derived PICALM-AU1 induced EndMT in PMVECs and enhanced angiogenesis in HPS.

PICALM-AU1 is a novel lncRNA. It has two exons with 368 bp in its coding sequence. Tissues exhibit low expression of PICALM-AU1 under normal physiological conditions. However, we detected the overexpression of PICALM-AU1 in the liver and lungs of HPS rats. Using immunopurification of primary cholangiocytes, laser capture microdissection, and FISH-immunohistochemistry, we also revealed that cholangiocytes were the primary source of hepatic PICALM-AU1 under physiologic and cholestatic conditions.

Exosomal PICALM-AU1 has a critical role in pathological angiogenesis in the lungs of HPS rats. Exosomes are small extracellular vehicles released by various types of cells, which can carry multiple cargos, including protein, DNA, mRNA, lncRNA and lipids [39-41]. Recently, exosomes have drawn significant attention as essential mediators of intercellular communication under physiological and pathological conditions [42]. Exosomal cargo in the serum, saliva, and urine can be used as potential biomarkers for low-invasive diagnoses of cancers [43-46]. Advanced stages of liver disease are irreversible and life-threatening and diagnosis of chronic liver disease at an early stage is challenging owing to the lack of non-invasive approaches and long asymptomatic disease progression. Several exosomal miRNAs and lncRNAs have been identified as potential diagnostic biomarkers for various liver diseases, including viral hepatitis, drug-induced liver injury, alcoholic liver disease, non-alcoholic fatty liver disease, hepatocellular carcinoma, and cholangiocarcinoma. In this study, we found that serum levels of exosomal PICALM-AU1 were closely correlated with hepatic PICALM-AU1 expression and severity of HPS. Data from patients with HPS showed that serum exosomal PICALM-AU1 is a potential diagnostic biomarker for HPS (Fig. S5A, B). However, it is unknown if functional PICALM-AU1 only resides in the liver. It is important to note the regulatory role of local PICALM-AU1 on EndMT in PMVECs.

EndMT is a core contributor of the formation of pulmonary microvasculature and component of pathways associated with the development of HPS. EndMT is reversible [47, 48]. Thus, it is important to understand the mechanisms involved in regulating EndMT for vascular remodeling and identification of novel therapeutic strategies aimed at reversing vascular remodeling to relieve the symptoms of HPS. This study focuses on the characteristics and functions of EndMT in vascular remodeling in HPS.

## Conclusions

In summary, we have demonstrated the roles of PICALM-AU1 in regulating EndMT during pathological blood vessel remodeling in HPS via exosome-mediated communication between distant organs. These findings highlight the clinical significance of Exo-PICALM-AU1 signaling as part of a novel therapeutic intervention in patients with severe liver injuries. The limitation of this work is we only detected PICALM expression differences in HPS patient serum exosomes, but there were no more data on PICALM expression location, differences expression in the HPS patients liver, and the function of exosomal PICALM-AU1 in the lung. And how to effectively intervene PICALM-AU1 to achieve the therapeutic effect of HPS patients. These unresolved issues are also the direction of the team’s efforts in the later stage.

## Conflict of interest

The authors declare that they have no conflict of interest.

## Acknowledgments

This project was supported by the National Science Foundation of China (81800060, 81671961, 81870422), China Postdoctoral Science Foundation (2017M623370) and Natural Science Foundation of Chongqing, China (xm2017085, cstc2020jcyj-msxmX0361).

## Author contributions

C.W.Y and K.Z.L designed the project, C.W.Y and Y.H.Y performed experiments, Y.C performed bioinformatics analyses, J.H and Y.J.L provided assistance with data analysis and curation, H.Y.Z and X.T collected HPS patients, J.L.N and J.T.G made animal model. B.Y and K.Z.L provided funding and supervision, C.W.Y and K. B wrote the manuscript, X.B.W and Z.Y.X revised the manuscript with input from all authors.

## List of Abbreviations

HPS: Hepatopulmonary syndrome
CBDL: common bile duct ligation
RNA: long noncoding
PICALM-AU1
miR144-3p
ZEB1

---

**Table 1.** Basic feature of HPS patients.

| | NO HPS<br>(N=73) | HPS (N=56) | $\chi^2$ | P value |
| --- | --- | --- | --- | --- |
| Age at diagnosis, median (min, max), y | 44 (18, 75) | 51.5 (21, 75) |  |  |
| Sex, male (female) [ratio] | 55 (18)<br>[75.3%] | 44 (12.0)<br>[78.6%] |  |  |
| median (min, max), cm | 168 (148, 177) | 165 (150, 180) |  |  |
| median (min, max), kg | 61 (39, 90) | 64.5 (42, 96) |  |  |
| pathology, (hepatitis B/others) | 45/28 | 43/13 | 3.351 | 0.086 |
| Aspartate transaminase, AST, U/L | 94.09±89.1 | 114.31±84.74 |  | 0.913 |
| Cerealthirdtransaminase, ALT, U/L | 144.35±190.59 | 88.91±78.2 |  | 0.001 |
| Albumin, mg/dl | 35.06±4.91 | 30.74±4.8 |  | 0.867 |
| Globulin, GLB, mg/dl | 30.43±7.3 | 31.53±7.84 |  | 0.237 |
| prothrombin time, PT | 13.98±3.5 | 18.77±5.07 |  | 0.003 |
| Meld index | 61.94±6.38 | 67.56±7.11 |  | 0.22 |
| Vertical dyspnea, (Yes/No) | 18/55 | 40/16 | 28.014 | <0.0001 |
| partial pressure of oxygen in artery | 98.56±2.65 | 86.92±19.11 |  | 0.027 |
| partial pressure of carbon dioxide in artery | 38.86±2.67 | 35.97±4.12 |  | 0.001 |
| alveolar-arterial oxygen tension difference, P(A-a)O<sub>2</sub>, mmHg | 8.62±6.91 | 28.56±8.52 |  | 0.023 |
| Type-B ultrasonic, (positive/negative) | 16/57 | 56/0 | 70.891 | <0.0001 |
| Clubbing digits | 27/46 | 23/33 | 5.13 | 0.024 |
| Spider angioma | 7.03±4.68 | 5.2±3.82 | 25.745 | 0.041 |

## Supplementary Figures

**Fig. S1.**
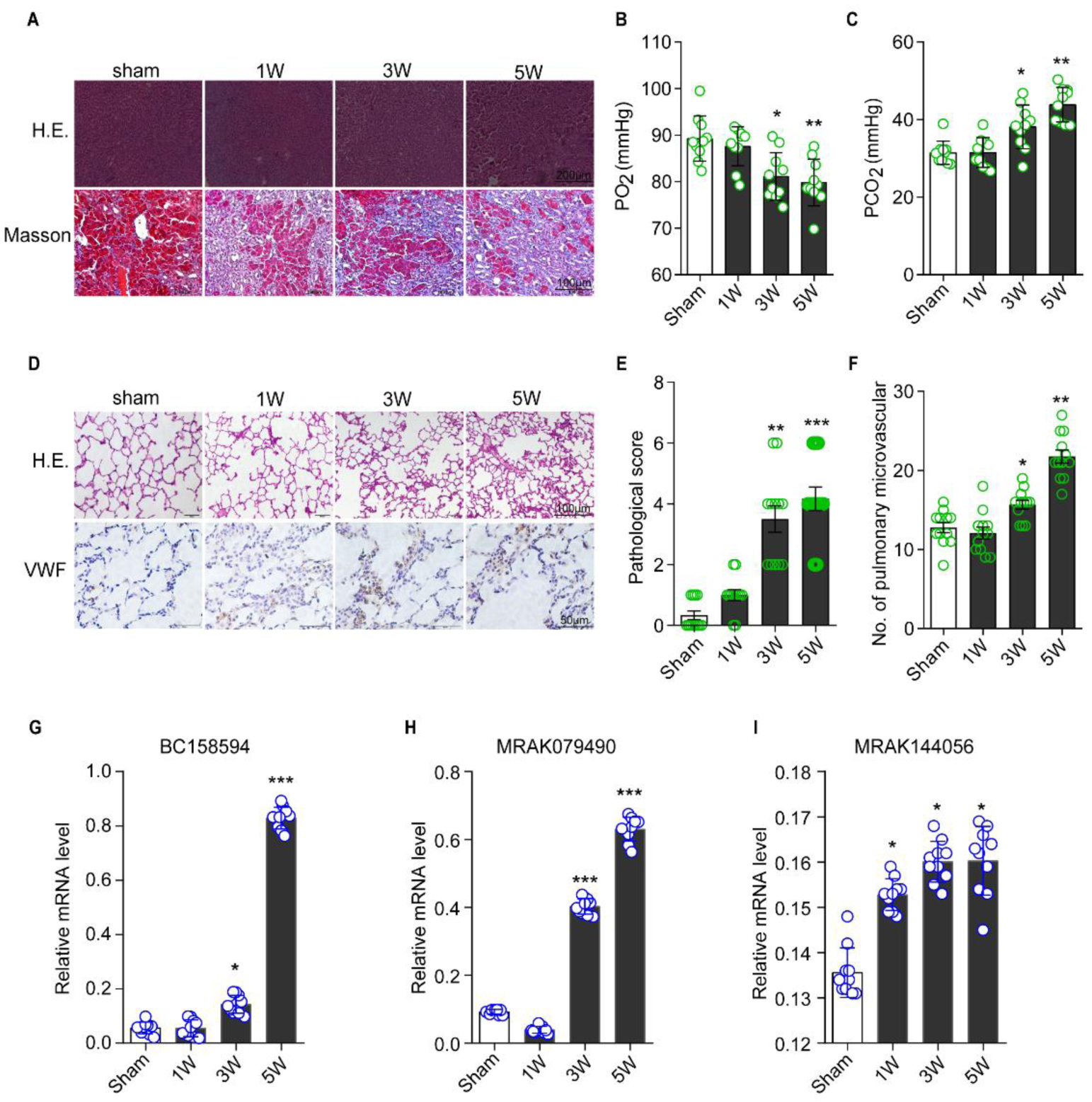
Generation of the rat model of CBDL. A, H.E. staining and Masson staining of the liver of HPS rats. B-C, PO_2_ and PCO_2_ in HPS rats. D, H.E. staining and lung vascular staining. E-F, Pathological score and the number of pulmonary microvasculature in HPS rat lung. G-I, qPCR for BC158594, MRAK079490 and MRAK144056 levels in the HPS liver.

**Fig. S2.**
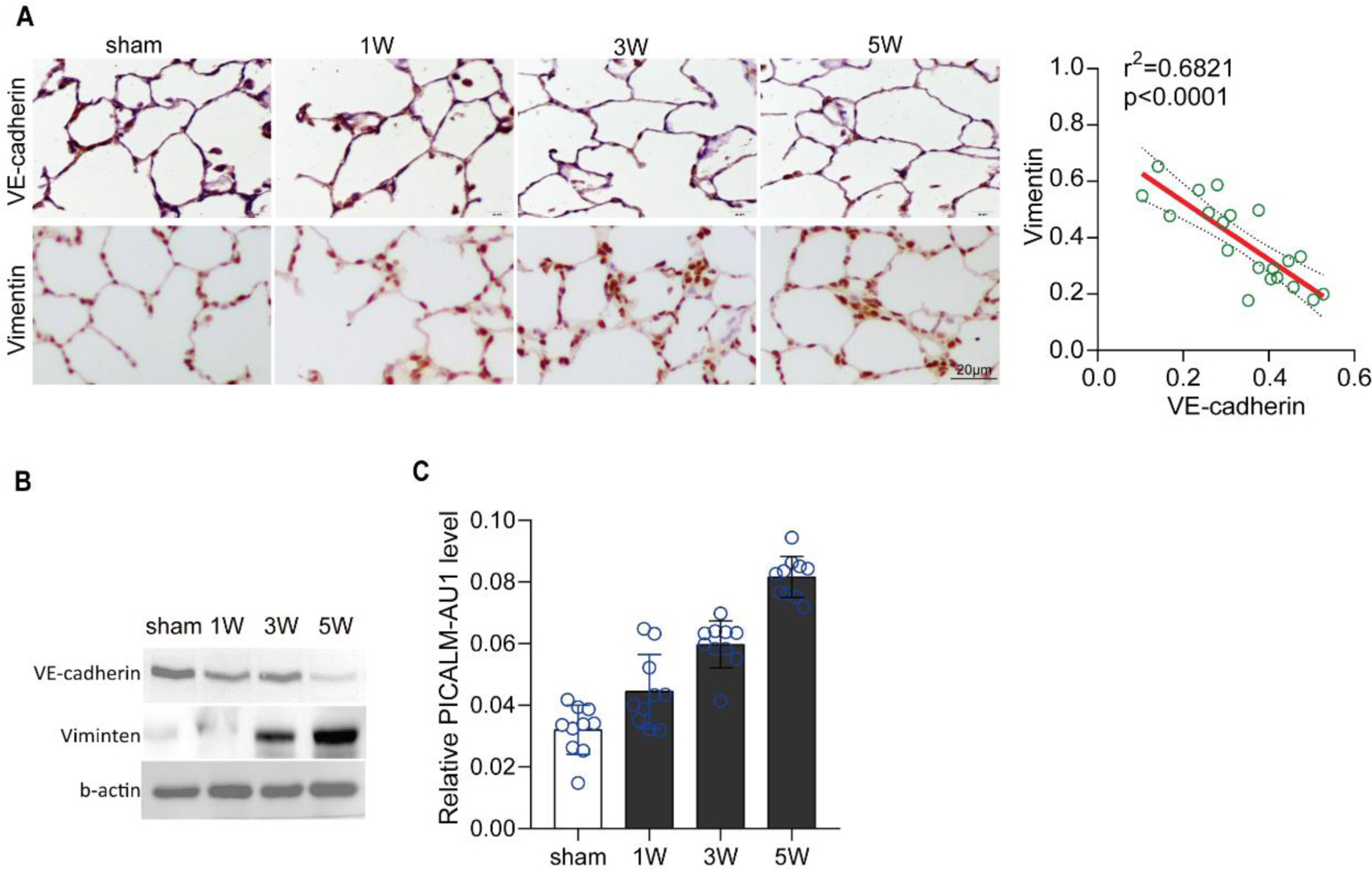
EndMT in PMVECs of the lungs of HPS rats. A, (Left) Immunofluorescence for the expression of VE-cadherin and vimentin in the lungs of CBDL rats. (Right) Linear regression analysis of vimentin and VE-cadherin expression. B, Western blotting for the protein levels of VE-cadherin and vimentin in the lungs of CBDL rats. β-actin was the internal control.

**Fig. S3.**
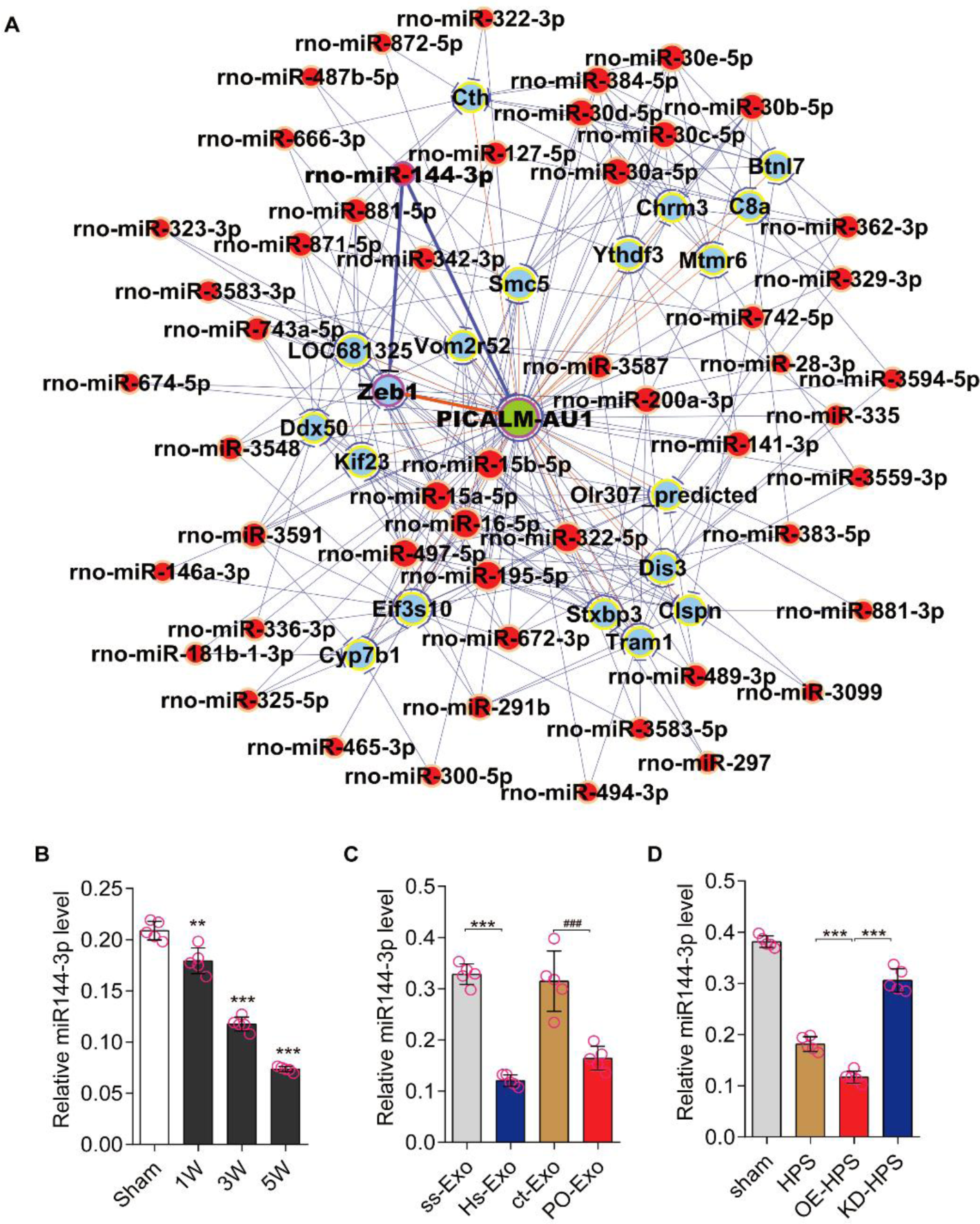
Regulatory network of PICALM-AU1. A, Regulatory network of PICALM-AU1. Genes colored in blue represent mRNAs and lncRNAs; those in red represent miRNAs. B, qPCR analysis miR144-3p levels in the lungs of HPS rats. Statistical significance relative to the sham group, *P<0.05, **P<0.01, ***P<0.001. C, qPCR analysis for miR144-3p levels in exosome-treated rats. Statistical significance relative to the ss-Exo treated group, *P<0.05, **P<0.01, ***P<0.001; relative to ct-Exo treated group, #P<0.05, ##P<0.01, ###P<0.001. D, qPCR analysis for miR144-3p levels in lentivirus-mediated PICALM-AU1-overexpressing or -depleted lungs of HPS rats. Statistical significance relative to the sham group, *P<0.05, **P<0.01, ***P<0.001; relative to the HPS group, #P<0.05, ##P<0.01, ###P<0.001.

**Fig. S4.**
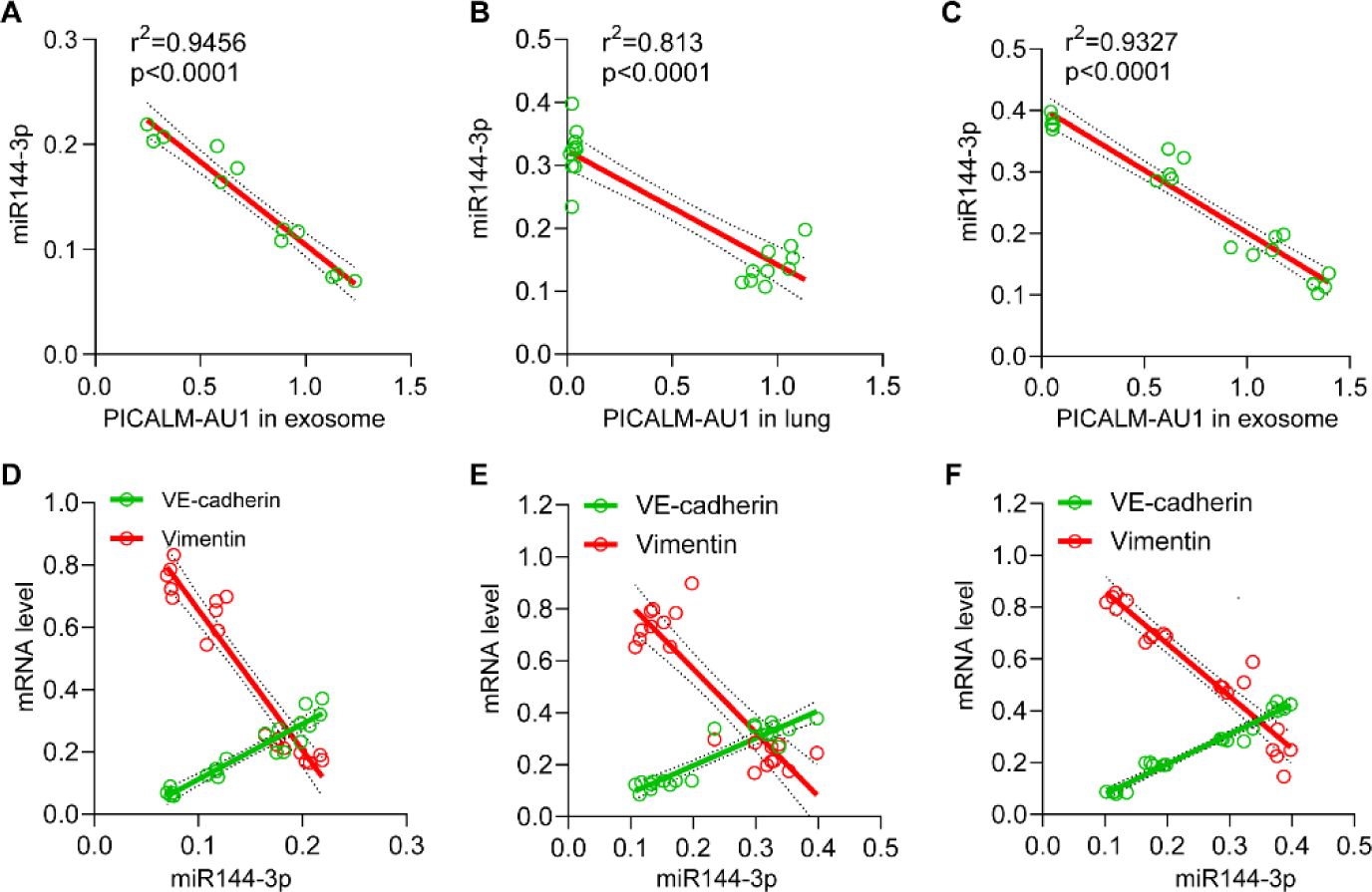
Correlation between the expression of miR144-3p and PICALM-AU1 and EndMT marker genes in HPS rats and PMVECs. A–C, Linear regression analysis for miR144-3p levels and PICALM-AU1 levels in the lungs. (A, in 3-wk-old CBDL rat model; B, in exosome-treated rat model; C, in PICALM-AU1-overexpressing or -depleted rat model) D–F, Linear regression analysis for the expression of miR144-3p and EndMT markers (VE-cadherin and vimentin) in the lungs. (A, in 3-wk-old CBDL rat model; B, in exosome-treated rat model; C, in PICALM-AU1-overexpressing or -depleted rat model)

**Figure.S5.**
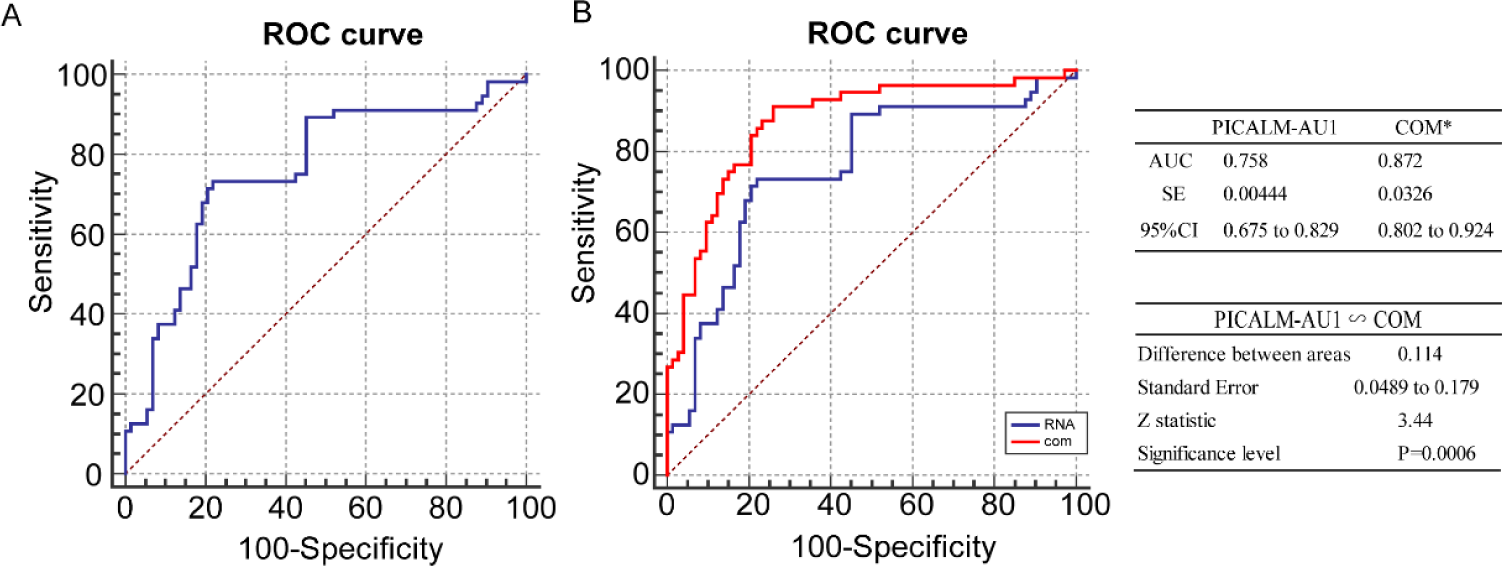
PICALM-AU1 level in HPS patient serum can be as a biomarker for HPS detection. A, ROC curve of PICALM-AU1; B, ROC curve of PICALM-AU1 and combine with spider angioma, total bile acid and vertical dyspnea.

**Table S1.**
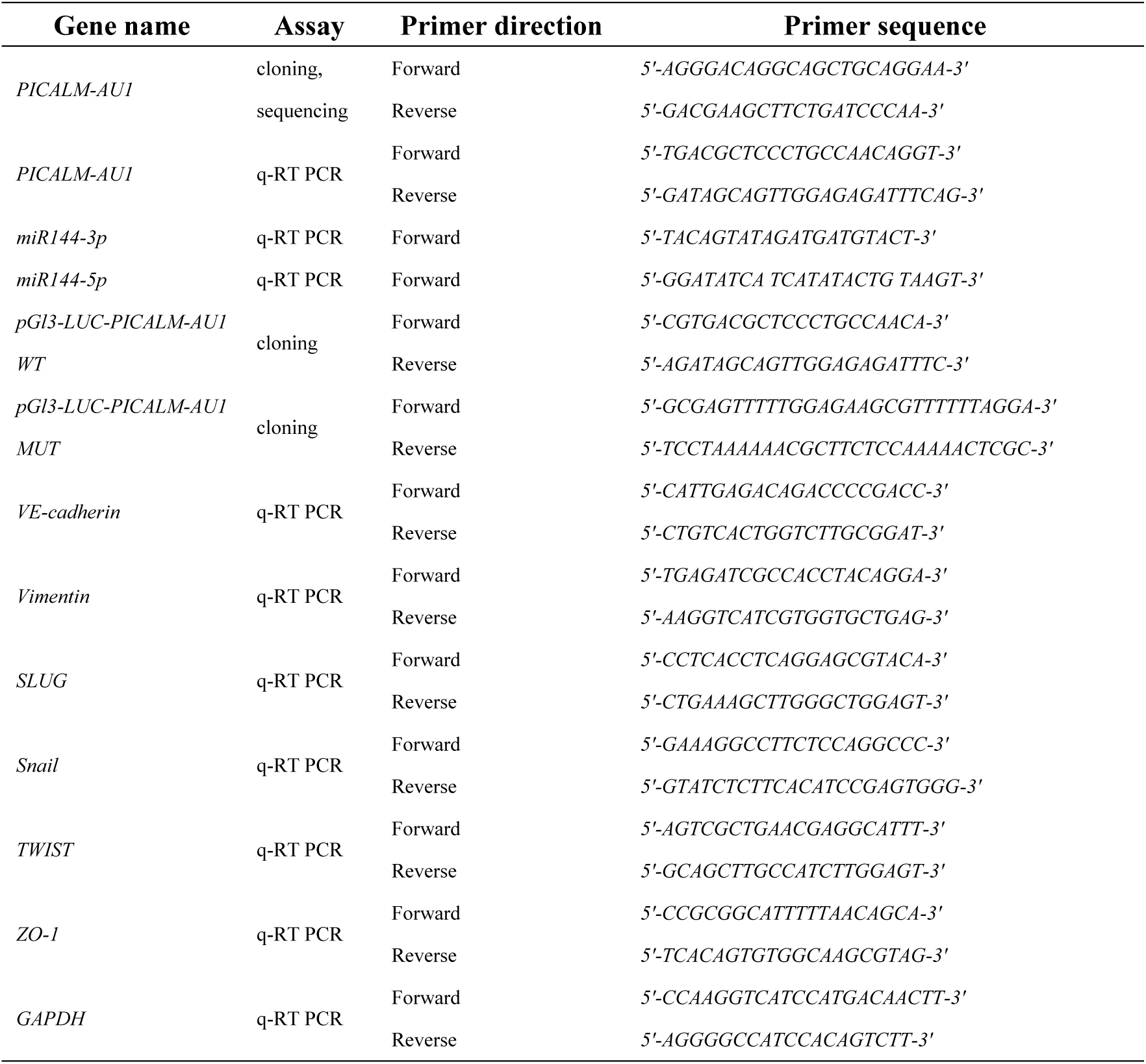
Primers for this study.

**Table. S2.** Antibodies information.

| Protein name | Assay | Manufacturer | NO. | Additional information |
| --- | --- | --- | --- | --- |
| VE-cadherin | Western blot, IHC, IF | abcam | ab33168 | Western blot, 1:2000; IHC and IF, 1:200 |
| Vimentin | Western blot, IHC | Cell Signaling Technology | 5741 | Western blot, 1:5000; IHC and IF, 1:200 |
| VWF | Western blot, IHC | abcam | ab6994 | Western blot, 1:5000; IHC and IF, 1:200 |
| CK19 | Western blot, IF | abcam | ab7754 | IHC and IF, 1:250 |
| CD63 | Western blot, IF | abcam | ab217345 | Western blot, 1:5000; IF, 1:200 |
| CD81 | Western blot | abcam | ab243887 | Western blot, 1:2000 |
| ZO-1 | Western blot, IF | abcam | ab96587 | Western blot, 1:2000; IF, 1:200 |
| ZEB1 | Western blot, IF | abcam | ab203829 | Western blot, 1:2000; IF, 1:100 |
| TWIST | Western blot | abcam | ab50581 | Western blot, 1:2000 |
| Snail | Western blot | abcam | ab229701 | Western blot, 1:2000 |
| SLUG | Western blot | Cell Signaling Technology | 9585 | Western blot, 1:2000 |
| N-cadherin | Western blot | Cell Signaling Technology | 14215 | Western blot, 1:2000 |
| β-actin | Western blot | Cell Signaling Technology | 3700s | Western blot, 1:10000 |

**Table. S3.** Prediction of miR144-3p target

| ID | miRNA | Target | Validation methods |  |  |  |  | SUM | # of papers |
| --- | --- | --- | --- | --- | --- | --- | --- | --- | --- |
|  |  |  | Strong evidence |  |  | Less strong evidence |  |  |  |
|  |  |  | Reporter assay | Western blot | qPCR | micro array | NGS |  |  |
| MIRT053494 | hsa-miR-144-3p | ZFX | ✓ | ✓ | ✓ | ✓ | ✓ | 5 | 3 |
| MIRT438343 | hsa-miR-144-3p | MET | ✓ | ✓ | ✓ | ✓ | ✓ | 5 | 2 |
| MIRT437437 | hsa-miR-144-3p | EZH2 | ✓ | ✓ | ✓ |  | ✓ | 4 | 1 |
| MIRT005869 | hsa-miR-144-3p | NOTCH1 | ✓ | ✓ | ✓ |  | ✓ | 4 | 2 |
| MIRT543081 | hsa-miR-144-3p | APP | ✓ | ✓ | ✓ |  | ✓ | 3 | 2 |
| MIRT731796 | hsa-miR-144-3p | ZEB1 | ✓ | ✓ | ✓ |  |  | 3 | 1 |
| MIRT731798 | hsa-miR-144-3p | ZEB2 | ✓ | ✓ | ✓ |  |  | 3 | 1 |
| MIRT735303 | hsa-miR-144-5p | CCNE1 | ✓ |  | ✓ | ✓ |  | 3 | 1 |
| MIRT735304 | hsa-miR-144-5p | CCNE2 | ✓ |  | ✓ | ✓ |  | 3 | 1 |
| MIRT006872 | hsa-miR-144-3p | MTOR | ✓ | ✓ | ✓ |  |  | 3 | 3 |
| MIRT007190 | hsa-miR-144-3p | PTEN | ✓ | ✓ | ✓ |  |  | 3 | 1 |
| MIRT007310 | hsa-miR-144-3p | NFE2L2 | ✓ | ✓ | ✓ |  |  | 3 | 2 |
| MIRT054851 | hsa-miR-144-3p | TTN | ✓ | ✓ | ✓ |  |  | 3 | 1 |
| MIRT731688 | hsa-miR-144-3p | MAP3K8 | ✓ | ✓ | ✓ |  |  | 3 | 1 |
| MIRT732183 | hsa-miR-144-3p | PTGS2 | ✓ | ✓ | ✓ |  |  | 3 | 1 |
| MIRT732445 | hsa-miR-144-3p | Adamts1 | ✓ | ✓ | ✓ |  |  | 3 | 1 |
| MIRT733172 | hsa-miR-144-3p | TUG1 | ✓ | ✓ | ✓ |  |  | 2 | 1 |
| MIRT733522 | hsa-miR-144-3p | XIST | ✓ | ✓ | ✓ |  |  | 3 | 1 |
| MIRT733751 | hsa-miR-144-3p | PBX3 | ✓ | ✓ | ✓ |  |  | 3 | 1 |
| MIRT734389 | hsa-miR-144-5p | RUNX1 | ✓ | ✓ | ✓ |  |  | 3 | 1 |
| MIRT734437 | hsa-miR-144-5p | TGIF1 | ✓ | ✓ | ✓ |  |  | 3 | 1 |
| MIRT734527 | hsa-miR-144-3p | IRS1 | ✓ | ✓ | ✓ |  |  | 3 | 1 |
| MIRT735210 | hsa-miR-144-5p | ROCK1 | ✓ | ✓ | ✓ |  |  | 3 | 2 |

